# Investigation into the Phenological Patterns of Trees in Socorro Plateau, Goa

**DOI:** 10.1101/2022.11.14.516458

**Authors:** C. Sathya, M. Murugash

**Affiliations:** Department of Botany, Bishop Heber College (Autonomous), Tiruchirappalli, Tamil Nadu, India

**Author notes:** Corresponding Author: Murugash M.

**Keywords:** Phenology, trees, climate change

## Abstract

Phenology is defined as the observation of biological events or life cycle phases that occur due to changes in climate and season. Primarily, phenology of all organisms is influenced by temperature fluctuations and quick climate changes which in turn regulate a variety of environmental factors. When discussing about the effects of climate change on ecosystems, phenology is especially important as many animals depend on the phenology of plants for their survival. Analysing the phenological patterns of a particular area can provide us with critical points on how ecosystems are affected by climate change. In this study, we have analysed the patterns of phenology for several trees and tried to identify any idiosyncrasies which may arise as a result of climate change and other similar factors. A total of 75 individuals belonging to 15 genera or 17 species were documented in Socorro plateau, Goa and their phenological patterns such as flowering, fruiting and leaf cover were recorded and analysed. The phenological patterns of the individual trees were compared in relation to each other such as in-between flowering and fruiting of the same species. In addition to this, the species were compared with each other in terms of their rate of flowering/fruiting to observe their inter-connected variations. Prospects for future research were identified to further promote studies in conservation biology of flora and fauna in the area and in climate change to mediate its effects.

## Introduction

Phenology is defined as the observation of biological events or life cycle phases that occur due to changes in climate and season. Primarily, phenology of all organisms is influenced by temperature fluctuations and quick climate changes which in turn regulate a variety of environmental factors. One of the best ways to understand plant effects is to question the plants themselves (1). Moreover, It has been assumed by some authors that phenology is the most responsive aspect of nature to warming and is also the simplest to observe (2). Leafing, flowering, fruiting, germination, and seed dispersal are examples of life cycle events that help us comprehend the phenology of plant communities. It’s still a mystery why leafing, flowering, and fruiting proceed in such a specific order (3). And the cause of variation in life cycle events for each unique species has yet to be established.

Several decades of analysis show that temperature rise is the prominent factor since industrial times that drives phenological aspects in plants. Numerous physical changes that had been linked to this warming include the sea level rise, melting of glaciers, decrease in snow and ice cover, increased depth to permafrost, and changes in wind patterns, temperature and precipitation (4).

Such deterioration has far-reaching implications, especially in terms of species ecology, population biology, reproductive biology, and phenology. Phenology is particularly important since it provides us with essential information on how climate change affects ecosystems (5). The phenological patterns of numerous trees were explored in this work, which could aid in determining species drift and also in biodiversity conservation.

## Materials and methods

### Study Area

Socorro plateau (15.5603°N latitude and 73.8487°E longitude) is part of Socorro village which lies five kilometers to the South-East of Mapusa town in Bardez taluka of Goa. It was initially part of a much larger village, Serula which was divided into the villages of Salvador do Mundo, Penha da Franca (Britona), Pomburpa, and Socorro.

### Observations

A total of 75 individuals belonging to 17 tree species were observed. Observations were carried out over four months starting from December 2021 to March 2022. Observations of the species were recorded in observation sheets. Every phenomenon was tabulated as numerical data in reference to a percentage (for example 20% of flowering, 45% of mature fruits, etc…). Observations were carried out every week with the number of field visits per week varying between 2-4 times and constant changes in phenological responses were observed under the given criteria. The cycle of events considered involves flowering, fruiting, and leafing.

### Photography

Photographs were taken in the field using Sony Bridge Camera Model DSC-HX400V. Photo plates of few species were also added that give evidence for the observed life cycle events.

### Data Analysis

These observations of flowering, fruiting and leafing which were taken as numerical values in terms of percentage were approximate in relevance to the area covered by the given parameter. Data were processed using Microsoft Excel Spreadsheet and analyzed.

## Results and discussion

The species and their families have been represented in the tabulation. For all documented species binomial and author citation were checked thoroughly.

**Table 1:** Individual Plant Species And Families.

| S.No | Name of Species | Family |
| --- | --- | --- |
| 1 | *Catunaregam spinosa* (Thunb.) Tirveng. | Rubiaceae |
| 2 | *Memecylon umbellatum* Burm.f. | Melastomataceae |
| 3 | *Careya arborea* Roxb. | Lecythidaceae |
| 4 | *Senegalia chundra* (Roxb. ex Rottler) Maslin | Fabaceae |
| 5 | *Falconeria insignis* Royle | Euphorbiaceae |
| 6 | *Lannea coromandelica* (Houtt.) Merr. | Anacardiaceae |
| 7 | *Zanthoxylum rhetsa* (Roxb.) DC | Rutaceae |
| 8 | *Terminalia bellirica* (Gaertn.) Roxb. | Combretaceae |
| 9 | *Ziziphus xylopyrus* (Retz.) Willd | Rhamnaceae |
| 10 | *Bombax ceiba* L. | Malvaceae |
| 11 | *Terminalia paniculata* B.Heyne ex Roth | Combretaceae |
| 12 | *Alstonia scholaris* (L.) R.Br. | Apocynaceae |
| 13 | *Caryota urens* L. | Arecaceae |
| 14 | *Ficus microcarpa* L.f. | Moraceae |
| 15 | *Anacardium occidentale* L. | Anacardiaceae |
| 16 | *Mangifera indica* L. | Anacardiaceae |
| 17 | *Ficus arnottiana* (Miq.) Miq | Moraceae |

### Analysis of Average Rate of Flowering and Fruiting in All Species

When analysing the rate of flowering and fruiting in all species, the rate of all the individuals for each species were compiled and tabulated below:

**Table 2:** Average Rate of Flowering and Fruiting in All Species.

| S.No | Name of Species | Flowering Rate<br>(% per week) | Fruiting Rate<br>(% per week) |
| --- | --- | --- | --- |
| 1 | *Catunaregam spinosa* (Thunb.)<br>Tirveng. | 1.46 | 2.40 |
| 2 | *Memecylon umbellatum* Burm.f. | 20.26 | 3.80 |
| 3 | *Careya arborea* Roxb. | 5.81 | 0 |
| 4 | *Senegalia chundra* (Roxb. ex<br>Rottler) Maslin | 0 | 2.34 |
| 5 | *Falconeria insignis* Royle | 8.23 | 1.20 |
| 6 | *Lannea coromandelica* (Houtt.)<br>Merr. | 15.29 | 2.08 |
| 7 | *Zanthoxylum rhetsa* (Roxb.) DC | 0 | 1.04 |
| 8 | *Terminalia bellirica* (Gaertn.)<br>Roxb. | 0 | 6.88 |
| 9 | *Ziziphus xylopyrus* (Retz.) Willd | 0 | 1.91 |
| 10 | *Bombax ceiba* L. | 6.18 | 0 |
| 11 | *Terminalia paniculata* B.Heyne ex Roth | 0 | 5.59 |
| 12 | *Alstonia scholaris* (L.) R.Br. | 0 | 8.54 |
| 13 | *Caryota urens* L. | 0 | 4.90 |
| 14 | *Ficus microcarpa* L.f. | 0 | 5.05 |
| 15 | *Anacardium occidentale* L. | 29.90 | 9.58 |
| 16 | *Mangifera indica* L. | 21.04 | 6.98 |
| 17 | *Ficus arnottiana* (Miq.) Miq | 0 | 9.48 |

### Average Rate of Flowering in All Species

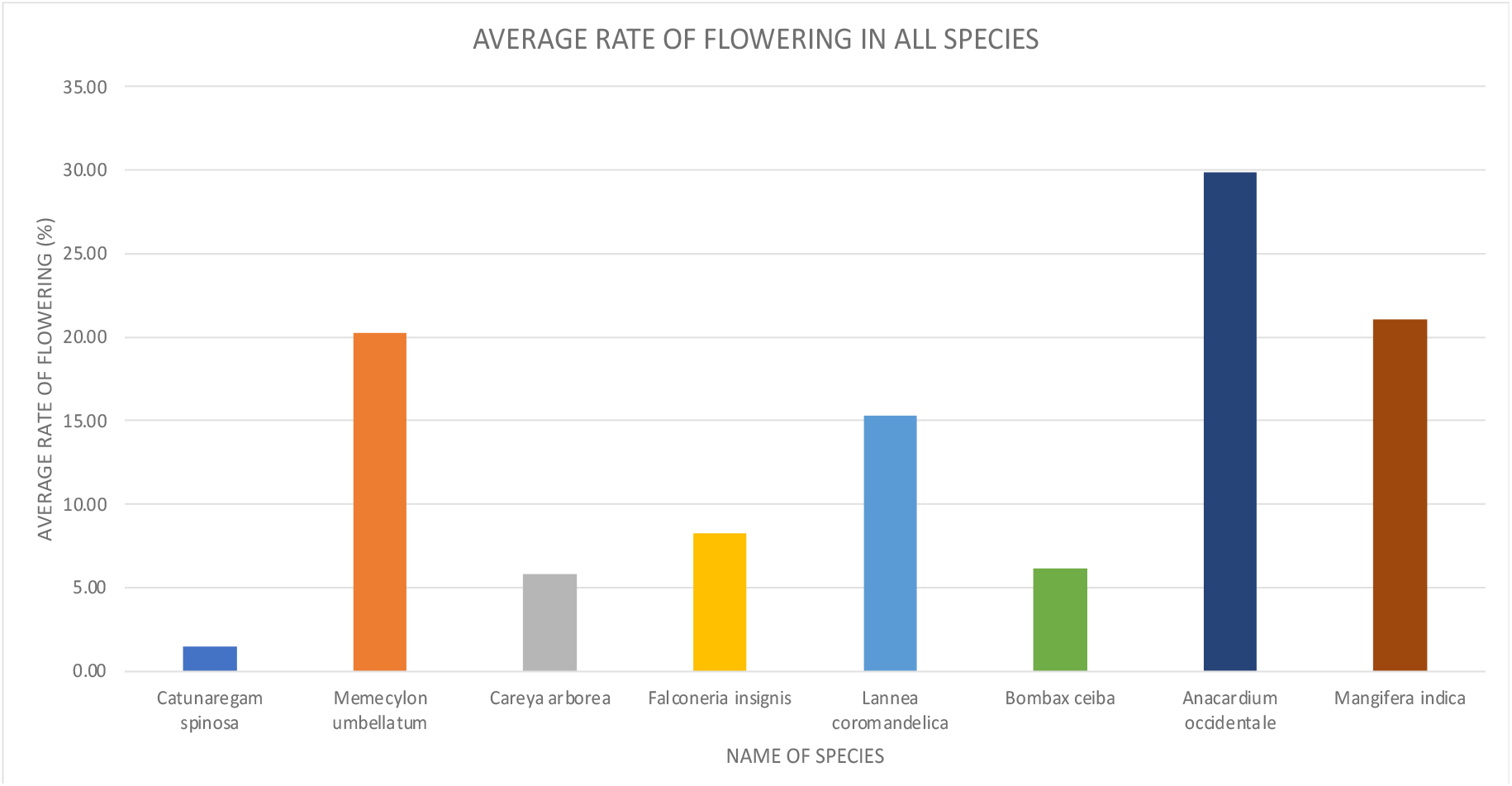

Analyzing the average rate of flowering in all species, expressed varying results. The species with the highest rate of flowering include *Anacardium occidentale, Mangifera indica* and *Memecylon umbellatum*. Among these, *A. occidentale* required special mention as there was an increased rate of flowering of about 30% every week while *M. indica* and *M. umbellatum* only increase by around 20%. Although the rate of flowering in *M. umbellatum* is comparatively less to species such as *A. occidentale*, it produces attractive violet blue inflorescence for longer durations and hence was a viable tool during the construction of botanical gardens and other floral landscapes (6); this is compounded by the fact that blue colour is rare in nature and provides beauty to a floral community (7). The species with the lowest rate of flowering include *Catunaregam spinosa, Careya arborea* and *Bombax ceiba*. Among these, *C. spinosa* has the lowest rate with about 1.4% increase in flowering per week while *C. arborea* and *B. ceiba* showed increased flower rate by approximately 6% every week. It should be noted that, even though *C. arborea* has one of the lowest rates, it is still early in its flowering cycle and so the flowering will possibly increase in the coming weeks of April and May. Also, in the case of both *C. arborea* and *B. ceiba*, increase in flowering is directly proportional to leaf-fall and this has been attested by previous studies (8,9). Therefore, by the time the individuals shown flowering, there was complete absence of leaf cover.

### Average Rate of Fruiting in All Species

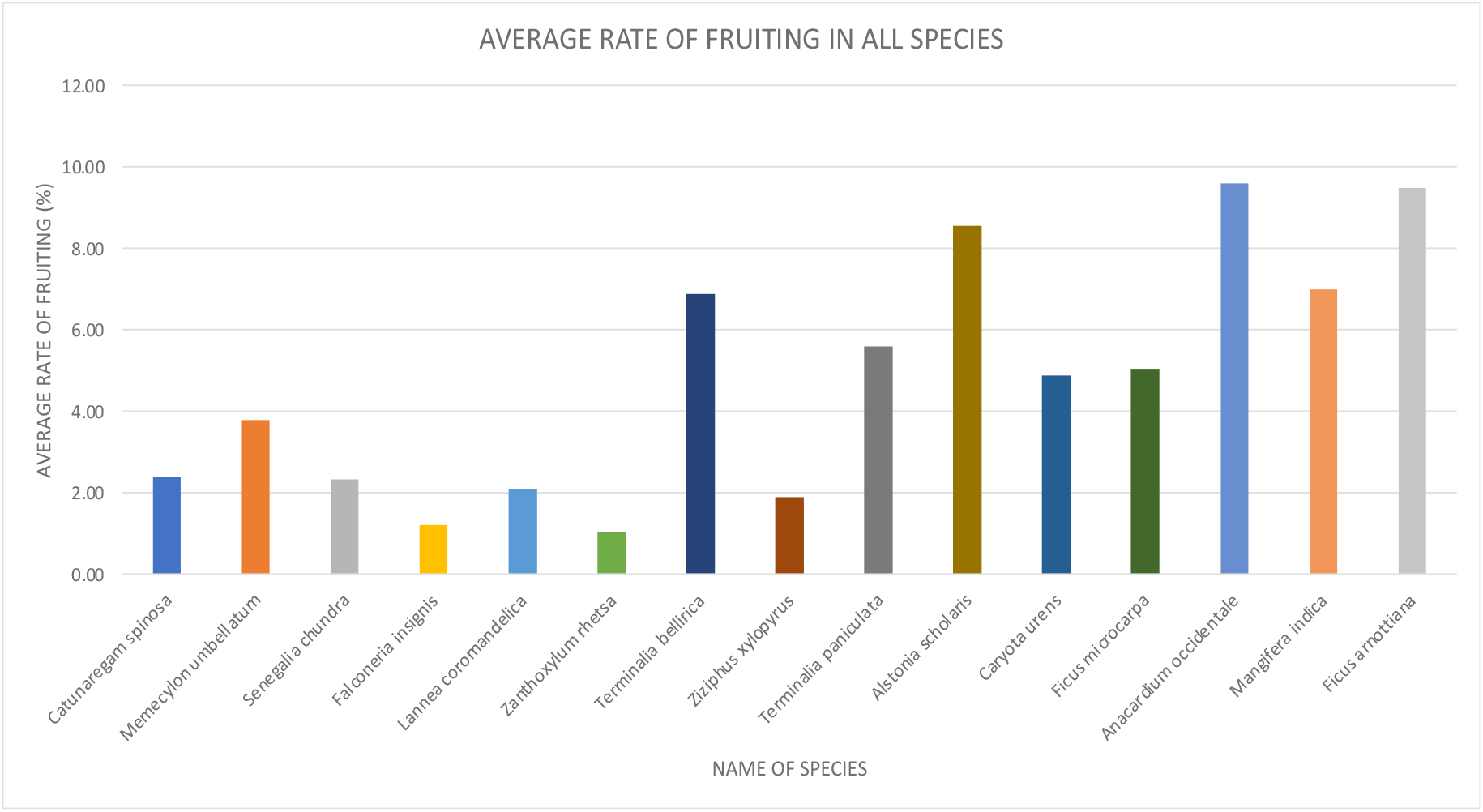

In this study, total of 15 species exhibited fruiting and so their average rate of fruiting have been compared. The overall rate of fruiting in all species is comparatively less to that of flowering as the maximum rate in fruiting was exhibited by *Anacardium occidentale* and *Ficus arnottiana* which achieve approximately 10% increase in fruiting per week while the maximum in flowering in nearly 30%. It is important to note that some species with the maximum rate of fruiting (*Anacardium occidentale and Ficus arnottiana*) are still undergoing fruiting and have yet to reach their peak while others such as *Alstonia scholaris* have already peaked and completed their fruiting season. In majority of the cases, with the few evergreen trees such as *Alstonia scholaris* and *Anacardium occidentale*, the reproductive events, be it flowering or fruiting were accompanied by leaf fall. Reproductive events, in general require large expenditure of energy and therefore occur during periods of low photosynthetic activity and/or high rates of reserve accumulation. Flowering, especially requires considerable amounts of energy to form non-photosynthetic tissue and nectar and so leaf fall maybe a possible adaptation by the plant to divert resources to flowering and also because it increases the visibility of flowers to pollinators (3,10). In relation to fruiting, in the tropics, the ripening of fruits generally coincide with the presence of numerous specialized frugivores which eat fruits throughout the year (3,11). The species with the lowest average rate of fruiting include *Falconeria insignis, Zanthoxylem rhetsa, Ziziphus xylopyrus* and *Lannea coromandelica*. *F. insignis* and *L. coromandelica* differ from the remaining two in the aspect both have just began their fruiting season and are expected to peak in the coming weeks in April/May.

### Variation in Phenological Patterns across Different Regions

The phenological patterns of a species in one climatic region may not be the same as in other regions. Therefore the consistencies in species across different regions were compared.

The phenological patterns of certain species such as *Memecylon umbellatum, Careya arborea, Lannea coromandelica* and *Caryota urens* were in line with previous studies

In *Memecylon umbellatum*, majority of the individuals of *Memecylon umbellatum* exhibited flowering and showed a peak in Week 4 (Mid-January) following this flowering remained stagnant for 3 weeks. After this sudden lag period, budding continued to increase from Week 6 (Early February) till the end of our study period. This corresponds with one of the four peaks revealed by Ratnayake and Yakandawala, (2006) in their study on the phenological patterns of *M. umbellatum* in Makandura, Sri Lanka. Similar to the study, we can also expect three more flowering peaks which will occur in April, May and June.

Canopy Cover in *Careya arborea* was a constant flux throughout the time period. Even though during the initial weeks, all individuals had a canopy cover of over 60%, the majority of the species showed a gradual decline from Week 6 (Early February) till the end of the study period. By the end of the study period, many individuals had canopy cover of less than 40% and this corresponded with gradual increase in flowering in the same timeframe. Hence, we may infer that the onset of flowering is corelated with leaf fall in *C. arborea*. Another study conducted in Assam, North India have reported similar findings although leaf fall begins sooner than flowering (8).

Flowering in *Lannea coromandelica* showed gradual increase from Week 3 (Mid-January). Budding does not show any significant peak while flowering exhibited gradual increase at Week 3 (Mid-January) and reached its peak in Week 11 (Mid-March). This is in line with previous studies conducted in the Vindhya Range of North India and in Rajasthan, India where flowering began in January and lasts till March before giving way for flowering (11,12).

Fruiting and Canopy cover of *Caryota urens* are constant for the entirety of the study period and this pattern is consistent with that of previous study conducted in Uttara Kannada district, Karnataka (13).

Although there are some species which are in line with previous studies irrespective of climatic conditions, there are some species are inconsistent with previous studies which include species such *as Falconeria insignis, Ziziphus xylopyrus, Alstonia scholaris* and *Anacardium occidentale*.

Flowering in *Falconeria insignis* occurs in all individuals except one. Budding shows gradual increase from Week 5 (Late January). It attained its highest peak in Week 11 (Mid-March) following which it shows sharp decline and this paves way for flowering. Although budding began in Week 5 (Late January), flowering occurs only 6 weeks later and so we can infer that the period of maturation of flowering in *F. insignis* is of approximately 48 days. However, in these individuals, the initiation of flowering is delayed by approximately 35 days as reported by previous study conducted in Uttara Kannada district, Karnataka where initiation of flowering usually occurs at the end of December followed by anthesis which occurred in the middle of January (13).

Fruiting in *Ziziphus xylopyrus* is in a gradual decline throughout the study period. Mature fruits especially, show a gradual decline for the majority of study period, but still show regular spike increases in percentage which correspond to the maturation of young fruits. Two peaks which occur in mature fruits were in Week 3 and Week 6 (Mid-January and Early February respectively). Peak in Week 6 (Early February) is followed by increase in ripening with peak in Week 7 (Mid-February). The ripened fruits eventually fall off and by Week 9 (Late February) no fruits remain. The ripening exhibited here is delayed by approximately a month when compared to other studies such as one by Bhat (1992) where ripening occurs in January itself.

Fruiting in *Alstonia scholaris* revealed a phase of steady decline throughout the majority of the study period ending by Week 9 (Late February). Mature fruits which start to appear before the beginning of study period reached its peak in Week 3 (Mid-January) showing a sharp decline from there with ripened fruits emerging only in Week 6 (Early February) and attaining peak in Week 7 (Mid-February) before declining soon after. Surprisingly this is accompanied in-between by the emergence of young fruits with a peak in Week 5 (Late January). This unnatural behavior of young fruits emerging after mature fruits maybe due to the persistence of mature fruits from the previous season and this leads to a large number of dried fruits at the end of this season. However, it is important to note that, there are another idiosyncrasy in the fruiting pattern of this plant which is that it is advanced by a period of 3-4 months as according to a previous study conducted in Assam, it usually showed fruiting in the period of May-July (14). Flowering period was varied among the different studies as in some it shows flowering between February-April and in others, it flowers between October-February (15,16).

Flowering in *Anacardium occidentale* seems to have begun before the study period and although high initially, first half of the study period expressed a gradual decrease remaining below 30% for the rest. This leads way for fruiting to occur. Flowering here, seems to be advanced by a period of 5 months where in another study conducted in Brazil, flower budding and emergence begins in May and continues till the first month of rainy season which is October (17).

## Conclusion

The various phenological patterns of flowering and fruiting were analyzed in Socorro plateau, Goa. Alongside analyzing the phenological patterns in the individual tree species in relation to each other, the average rate of flowering and fruiting in all the species were compared and analyzed. The phenological patterns of the current study were compared to that of previous studies. Certain species were found to be inconsistent and these inconsistencies maybe due to several reasons such as different environmental cues in different regions, availability of water, soil conditions, weather patterns etc. and hence may not be attributed to simply climate change due to the absence of long-term monitoring data. Future studies may entail studying the phenological patterns for long periods such as multiple years to assess the effect of climate change and also the effect, various fauna such as birds, mammals and insects have on the phenology of plants and vice versa.

## Supporting information

Photo Plates

## Acknowledgement

We thank Dr. Malapati K. Janarthanam, Senior Professor, Department of Botany, Goa university for his supervision over this project and providing the necessary resources needed for the successful completion of this project.

