## Supplementary material for "Investigation into the Phenological Patterns of Trees in Socorro Plateau, Goa": Photo Plates

PLATE 1: *Alstonia scholaris* (L.) R.Br.

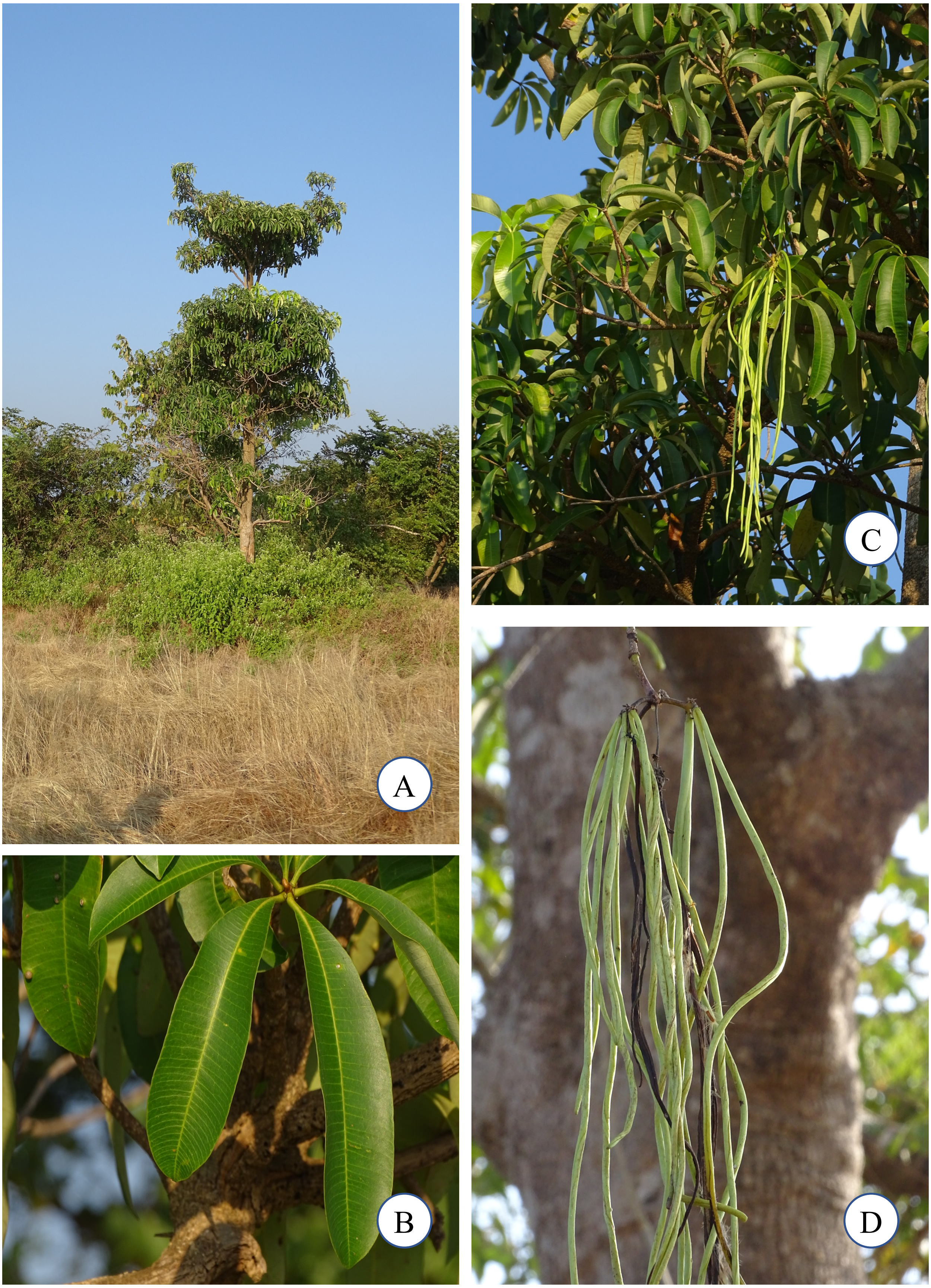


A. Habit; B. Leaves; C. Branches; D. Fruits

PLATE 2: *Anacardium occidentale* L.

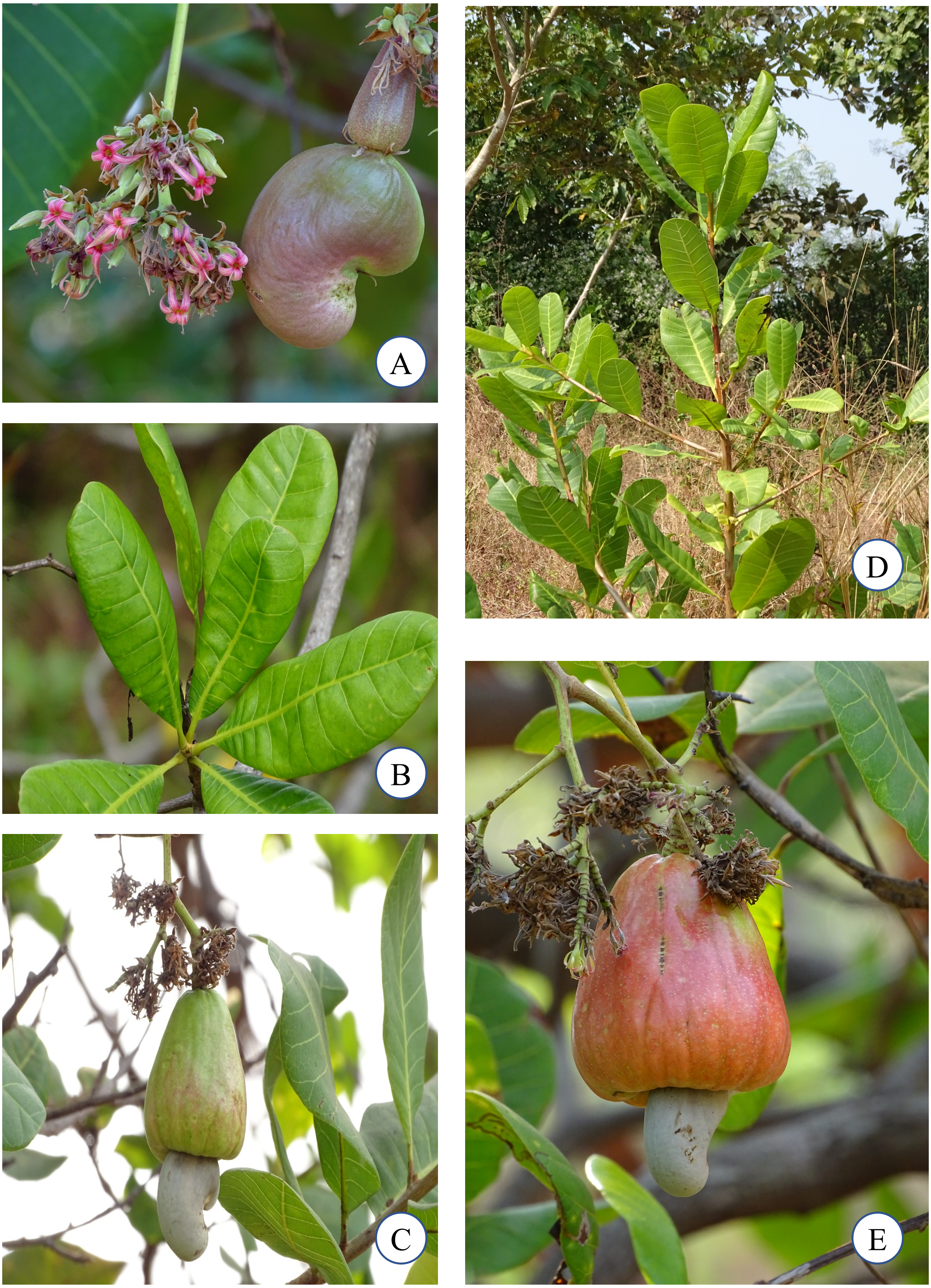


A. Flower and Young Fruit; B. Leaves; C. Mature Fruit; D. Young Tree; E. Ripened Fruit

PLATE 3: *Bombax ceiba* L.

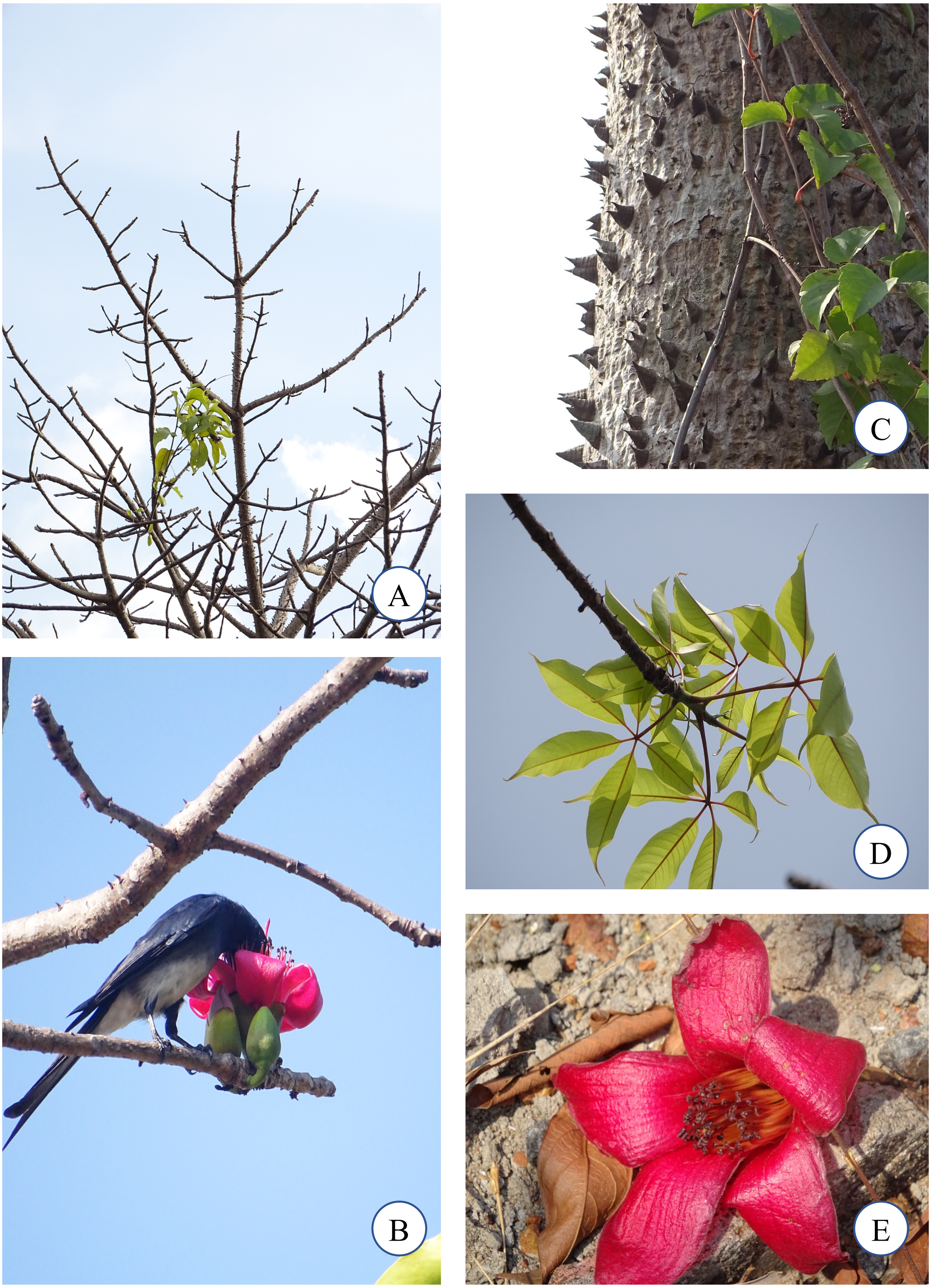


A. Habit; B. White-bellied Drongo (*Dicrurus caerulescens*) feeding on flower nectar; C. Bark; D. Leaves; E. Flower

PLATE 4: *Careya arborea* Roxb.

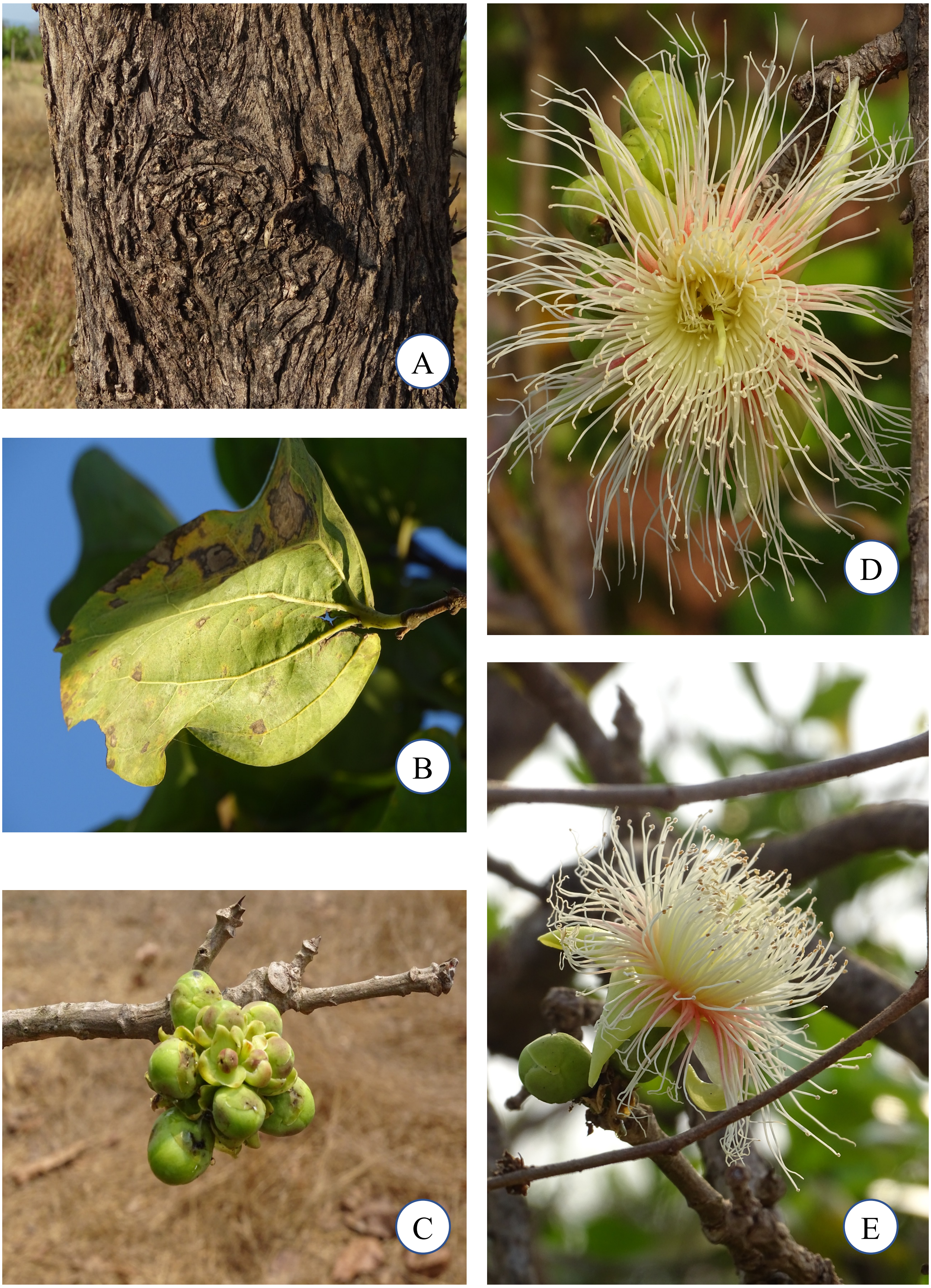


A. Bark; B. Leaf; C. Buds; D & E. Flowers

PLATE 5: *Catunaregam spinosa* (Thunb.) Tirveng.

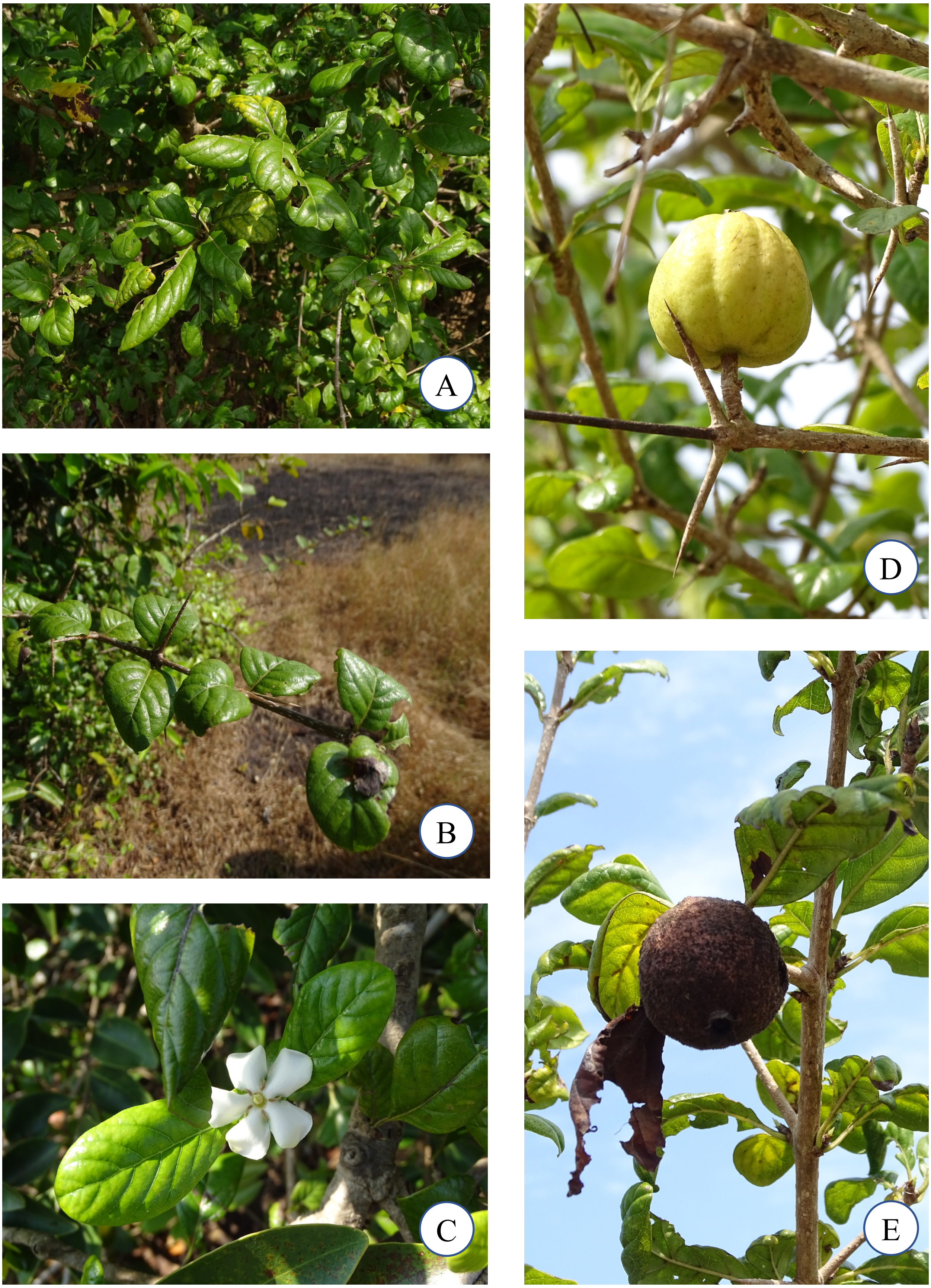


A & B. Branches; C. Flower; D. Mature Fruit; E. Dry Fruit

PLATE 6: *Falconeria insignis* Royle

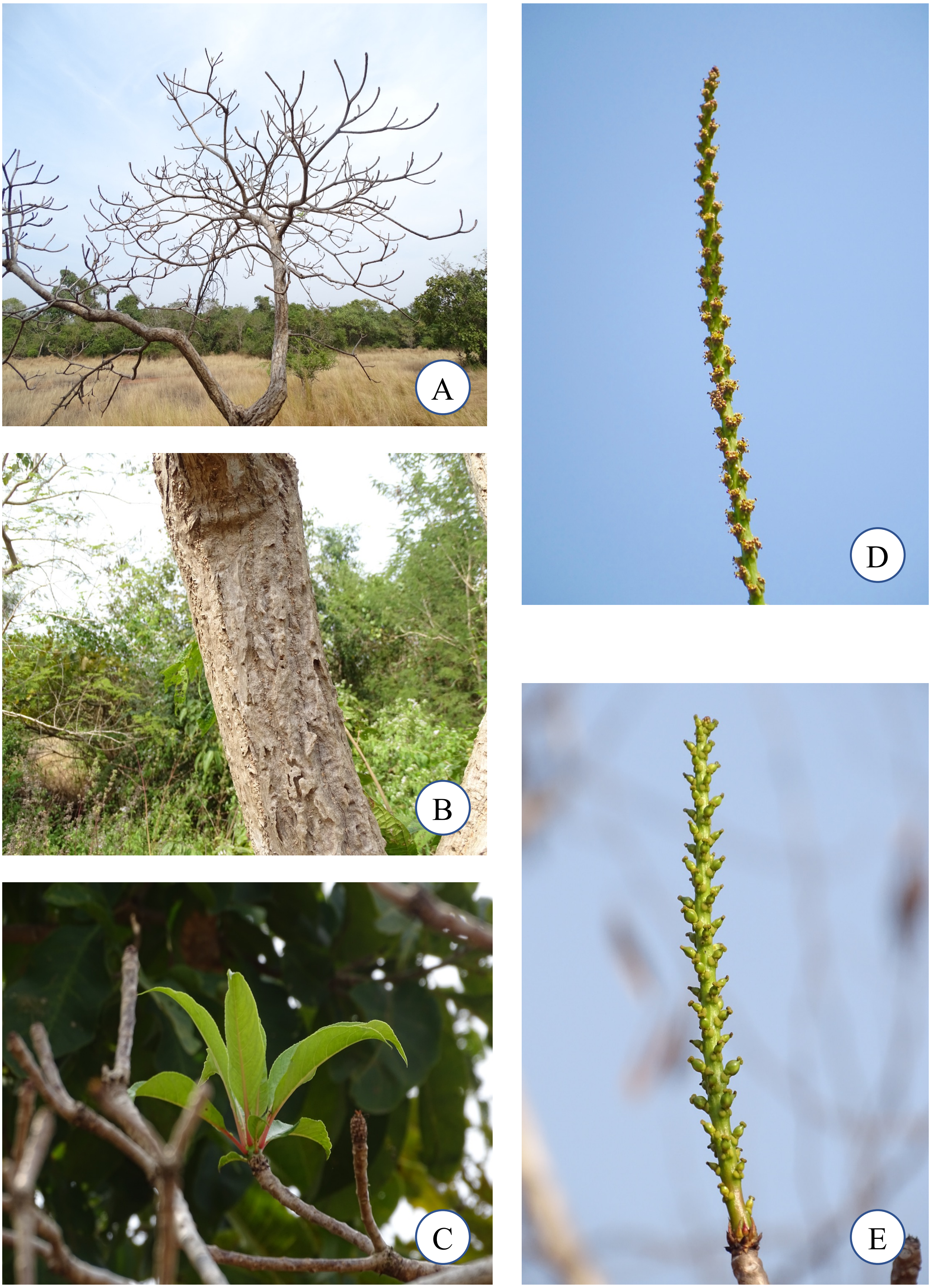


A. Habit; B. Bark; C. Leaves; D. Male Flower; E. Female Flower

PLATE 7: *Mangifera indica* L.

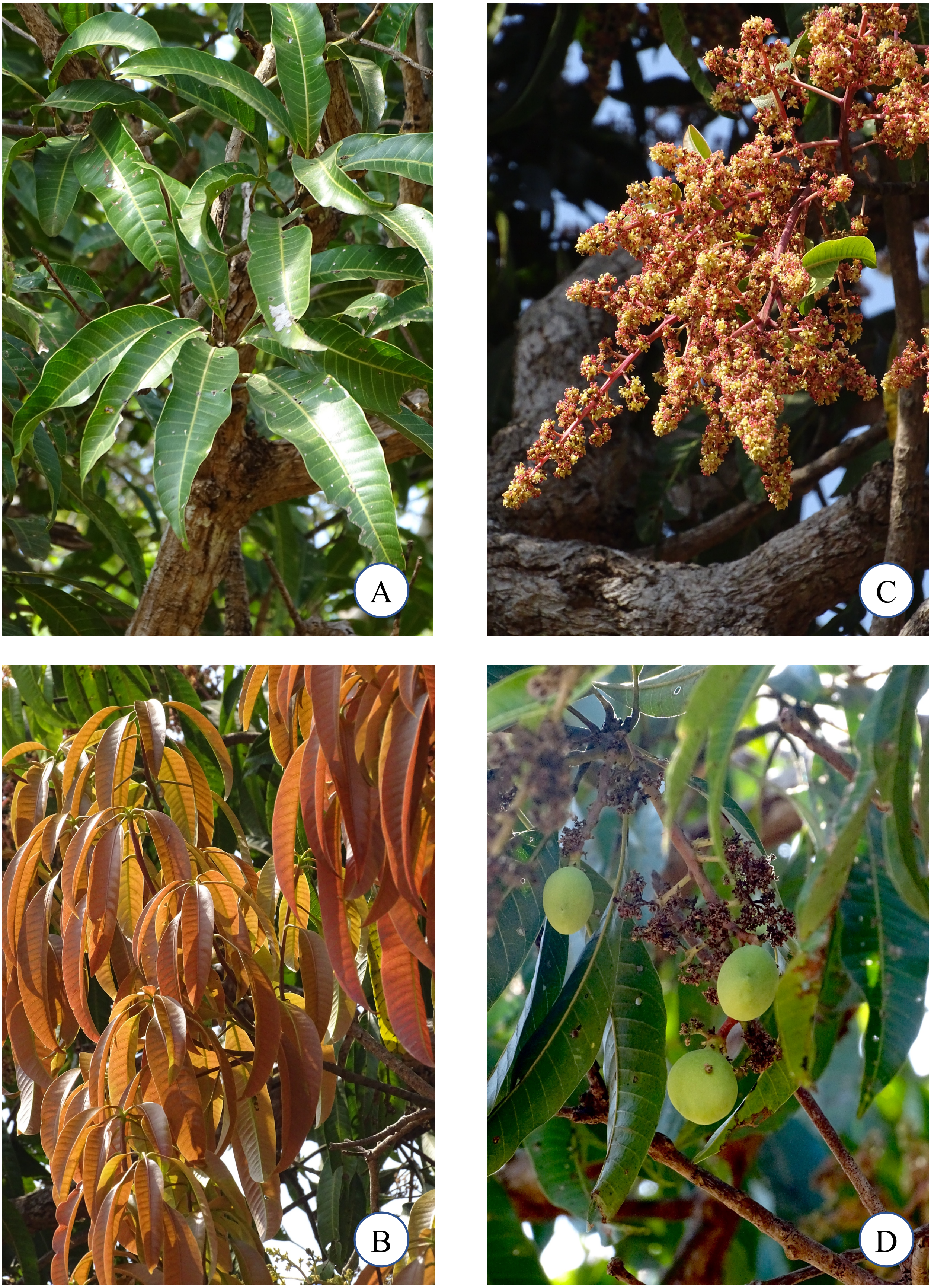


A & B. Leaves; C. Flowers; D. Young Fruits

PLATE 8: *Memecylon umbellatum* Burm. f.

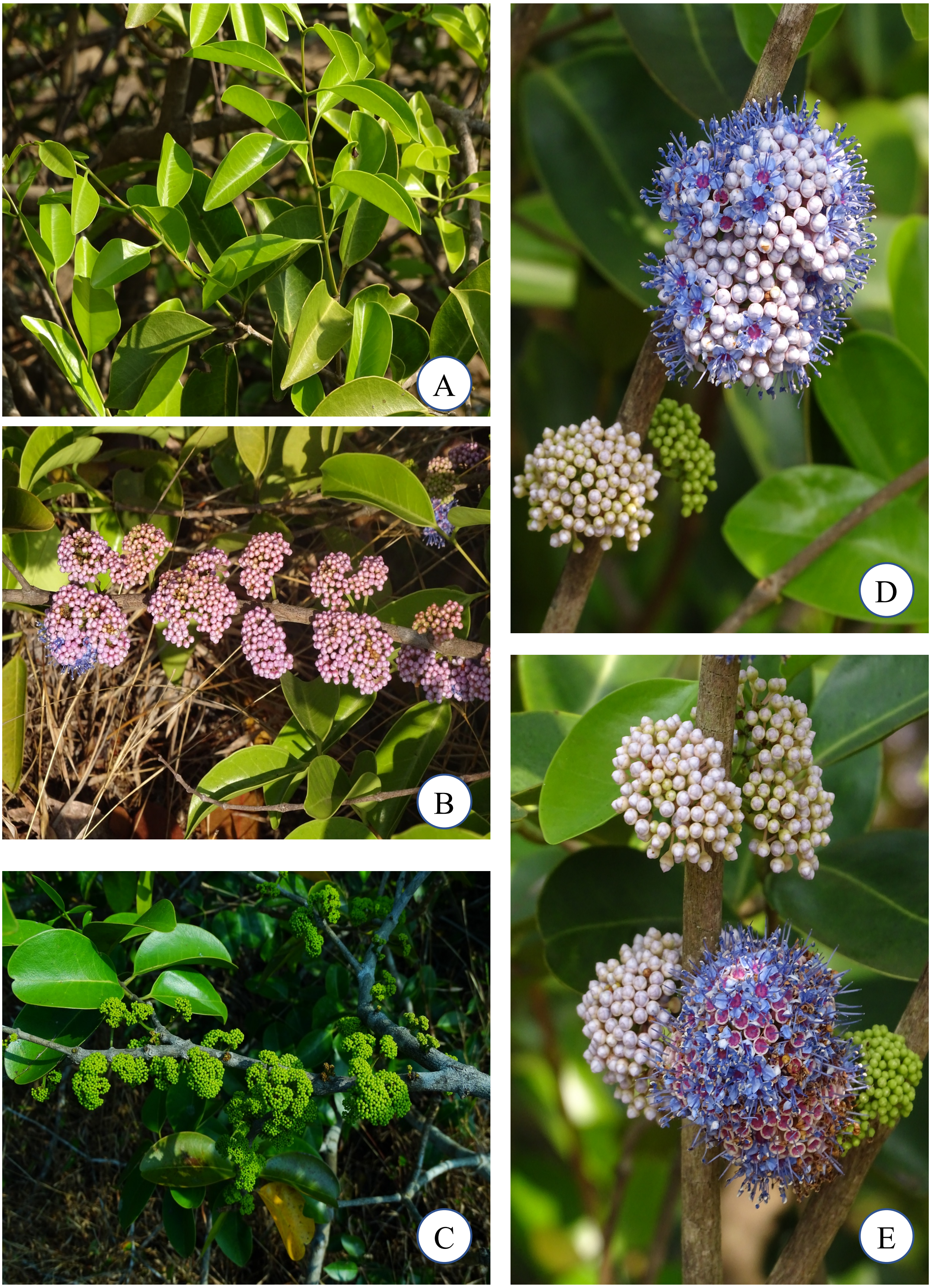


A. Leaves; B – E. Flowers at various stages of maturity
